# A role for the *MEGF6* gene in predisposition to osteoporosis

**DOI:** 10.1101/2020.01.09.900696

**Authors:** Craig C. Teerlink, Michael J Jurynec, Rolando Hernandez, Jeff Stevens, Dana C. Hughes, Cherie P. Brunker, Kerry Rowe, David J. Grunwald, Julio C. Facelli, Lisa A. Cannon-Albright

## Abstract

Osteoporosis is a common skeletal disorder characterized by deterioration of bone tissue in later life. The set of genetic factors contributing to osteoporosis is not completely specified. High-risk osteoporosis pedigrees were analyzed to identify genes that may confer susceptibility to disease. Candidate predisposition variants were identified initially by whole exome sequencing of affected-relative-pairs, approximately cousins, from ten pedigrees. Variants were filtered on the basis of population frequency, concordance between pairs of cousins, affecting a gene associated with osteoporosis, and likelihood to have functionally damaging, pathogenic consequences. Subsequently variants were tested for segregation in 68 additional relatives of the index carriers. A rare variant in *MEGF6* (rs755467862) showed strong evidence of segregation with the disease phenotype. Predicted protein folding indicated the variant (Cys200Tyr) may disrupt structure of an EGF-like calcium-binding domain of MEGF6. Functional analyses demonstrated that complete loss of the paralogous genes *megf6a* and *megf6b* in zebrafish resulted in significant delay of cartilage and bone formation. Segregation analyses, in-silico protein structure modeling, and functional assays support a role for *MEGF6* in predisposition to osteoporosis.

## INTRODUCTION

Osteoporosis is a common skeletal disorder characterized by deterioration of bone mineral density (BMD) and increased risk of fracture (1). Identification of genetic factors that have a strong influence on disease could enhance screening and knowledge concerning biological pathways involved. Genome-wide association studies (GWAS) have identified over 500 common single nucleotide polymorphisms (SNPs) associated with BMD (2), revealing several relevant biological pathways including genes involved in WNT signaling, the RANK/RANKL/OPG pathway, and genes involved in endochondral ossification. GWAS studies have revealed that osteoporosis is highly polygenic. SNPs identified thus far are estimated to account for about 20% of trait variation (3), whereas overall heritability estimates for BMD are higher, ranging from 50 to 90% (4, 5). The difference may in part be due to rare genetic variations whose effects are not typically detectable in GWAS. Family based designs present a complementary approach that may provide opportunities to discover rare, penetrant variants (6). Using a multi-generational pedigree resource, we performed exome sequencing on affected-relative-pairs with a deficiency in BMD from 10 high-risk osteoporosis pedigrees to identify rare, coding variants that may be contributing to osteoporosis risk in these families. Analysis of this resource identified a rare coding variant in the *MEGF6* (*multiple epidermal growth factor-like protein 6*, alternately known as *EGFL3*) gene that segregates to additional family members with osteoporosis. Computational and functional analyses indicate that the *MEGF6* variant likely disrupts wildtype (WT) protein function and that zebrafish paralogues contribute to bone and cartilage formation *in vivo*.

## RESULTS

### Variant prioritization

To identify genes associated with osteoporosis, multigenerational pedigrees were identified in which near-cousin pairs as well as additional individuals were confirmed to have a diagnosis of osteoporosis (see Materials and Methods). Exome sequencing of affected near-cousin pairs in each of ten pedigrees identified a total of ∼45K variants. Among these, 548 coding variants in 480 genes were rare (MAF<0.005 in EXAC browser (7)) and concordant between a sequenced pair of affected cousins. Restricting candidates to identity-by-descent (IBD) regions for sequenced pairs as identified by shared genomic segments (SGS) analysis (8) further limited the selection to 125 rare, shared candidate variants in 111 genes. A search of PubMed for each of the 111 genes to identify previous reported association to osteoporosis further limited selection to variants in 12 candidate genes. In a fourth filtering step, variants that were scored as ‘damaging’ by at least 8 of 10 in-silico pathogenicity prediction algorithms included in ANNOVAR variant annotations were retained, resulting in 3 candidate genes (*MEGF6* (rs755467862), *PAX8* (rs1269341622), and *UMPS* (rs777350640)).

### Confirmation of candidate variants via segregation

Six additional carriers of the *MEGF6* variant c.599 T>C; p.Cys200Tyr (rs755467862) were identified among 14 relatives of the index cousins that were assayed for the variant. Among the eight variant carriers identified in the pedigree, the sequenced index cousins as well as one relative had a physician-confirmed diagnosis of osteoporosis, four additional individuals had a physician confirmed diagnosis of osteopenia, and one individual was unaffected (screened in the 50th decade of life); four of the eight carriers had a medical history of bone fractures, and four had been treated previously for osteoporosis (Figure 1). Of the remaining six assayed relatives that did not carry the variant, none had a physician-confirmed diagnosis of osteoporosis. The de-identified pedigree segregating rs755467862 is depicted in Figure 1 and had significant evidence of segregation to affected relatives (osteoporosis or osteopenia, RVsharing p=0.0011) after correcting for 3 candidate variants (α=0.05/3=0.017) (9).

**Figure 1.**
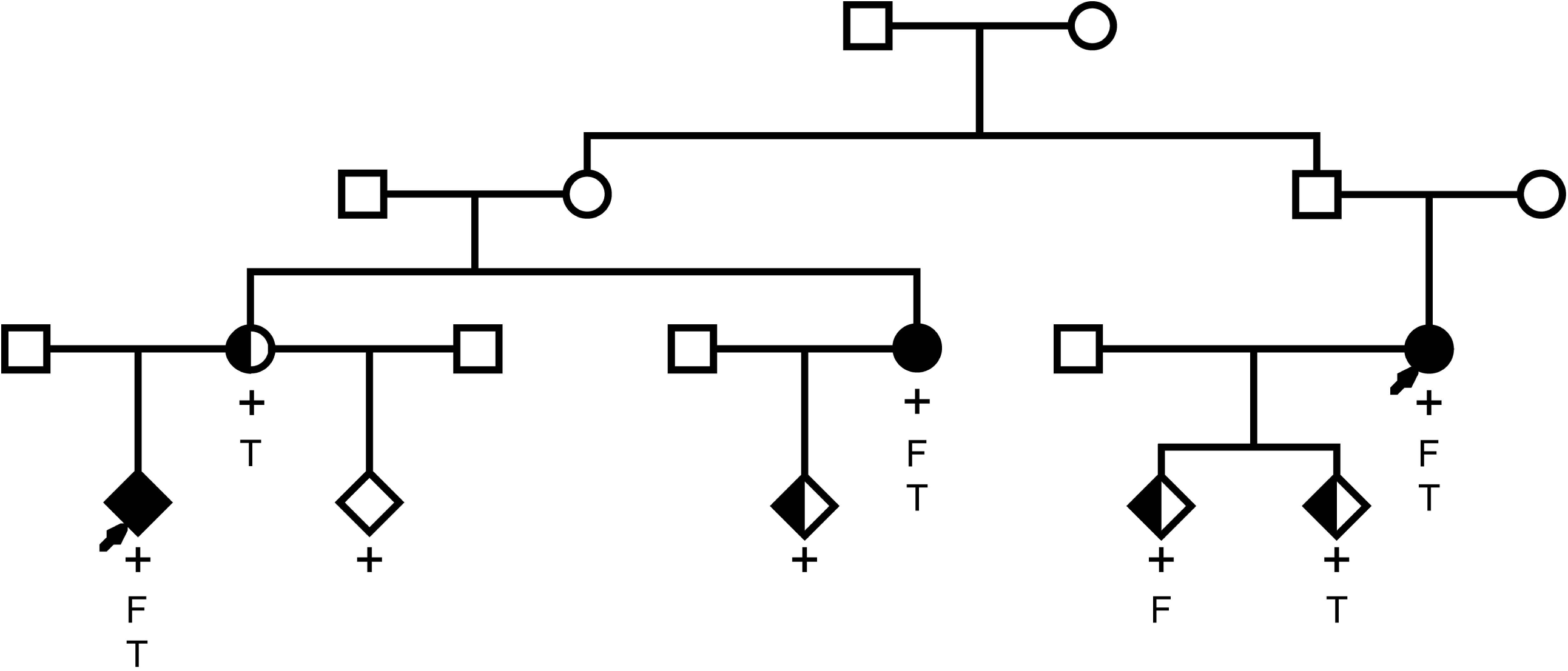
Osteoporosis pedigree segregating *MEGF6* variant. Physician confirmed diagnosis of osteoporosis is denoted by black fill and physician confirmed diagnosis of osteopenia is denoted by half-shading. Arrows indicate index sequenced subjects, ‘+’ indicates confirmed carriage of the variant, ‘F’ indicates multiple fractures in the patient’s medical history, and ‘T’ indicates previous treatment for osteoporosis.

### In silico functional evaluations of rs755467862

Variant rs755467862 is a non-synonymous variant substituting base T for C (c.599 T>C) in exon 5 that codes a Cysteine > Tyrosine amino acid substitution (p.Cys200Tyr). Population frequency has been estimated (MAF/number of alleles tested) as T=4e-5/243,578 (10). Variant rs755467862 was predicted to be damaging by almost all of the functional prediction software included in Annovar (11), including Polyphen (12), LRT_pred (13), MutationTaster (14), FATHMM (15), Radial_SVM (16), LR score (16), but not SIFT (17). The GERP score for the variant is 5.0, indicating strong evidence for conservation of the reference allele across species (18).

The candidate variant rs755467862 produces an amino acid substitution Cys200Tyr in the second EGF-like calcium binding domain of MEGF6. As MEGF6 is a large protein of 1541 amino acids, the entire protein is not amenable to existing 3D structure prediction methods. I-TASSER software (19) predicted the structure of the affected EGF-like calcium binding domain (residues 161-201) for both wild type and variant amino acid sequences, resulting in two distinct candidate structures for the wild type and five structures for variant proteins. From the ten possible comparisons, five (50%) showed substantial differences between the wild type and variant predicted structures. An example of this is presented in Figure 2 where it is apparent that the amino acid substitution disrupts a disulfide bond between residues 187 and 200 in the second EGF-like calcium-binding domain, where a helix is formed instead of an open loop. Similar behavior was observed in the other comparisons. The main structure of EGF-like domains is a two-stranded β-sheet followed by a loop where the calcium ion binds (20). The collapse of this loop may eliminate this calcium binding site in MEGF6, which may decrease, or even eliminate, the functionality of the protein.

**Figure 2.**
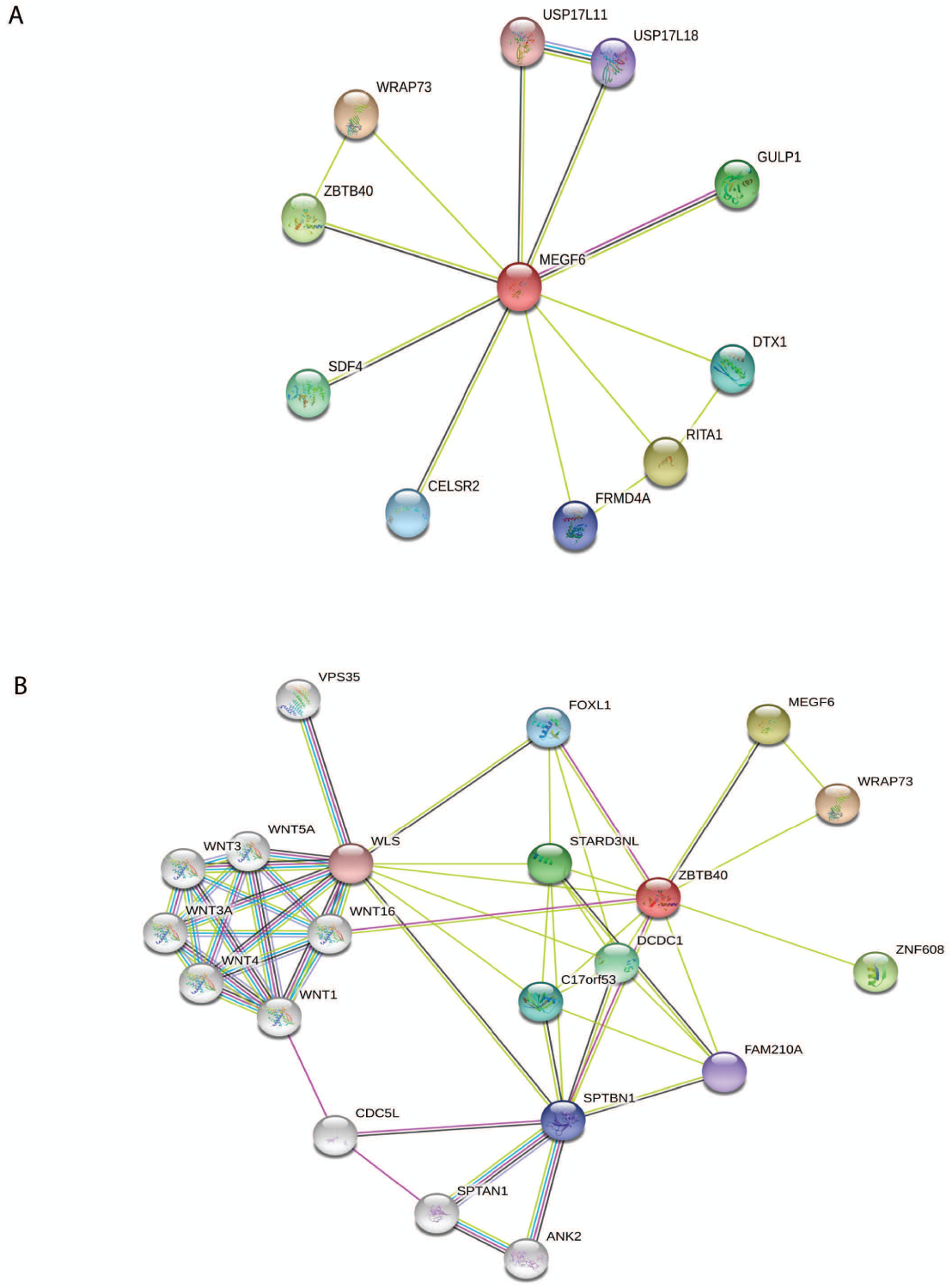
Comparison of the structure of *MEGF6* wild type (tan) and variant (blue) in the region of the Cys200Tyr. The disulfide bond between residues 187 and 200 have been highlighted in the wild type and mutant structures in red and yellow, respectively. The elimination of a loop can be visually observed, which is likely to adversely affect the calcium binding function of the first cbEGF-like domain in *MEGF6*.

### Protein interaction network

As variation in *MEGF6* expression or function has not been shown to be associated with bone development or homeostasis, we analyzed whether its protein product might have potential interactions with proteins more directly related to bone formation. Analysis with the STRING database tool (21) showed MEGF6 is related to pathways involved in various biological processes (Figure 3A) including angiogenesis (*EMILIN1*, a protein involved in vessel assembly (22), and most notably, osteoporosis (*ZBTB40* (23, 24)). Figure 3B contains a STRING graph re-centered on *ZBTB40* that shows connections with *WNT16*, which has been previously associated with BMD (25).

**Figure 3.**
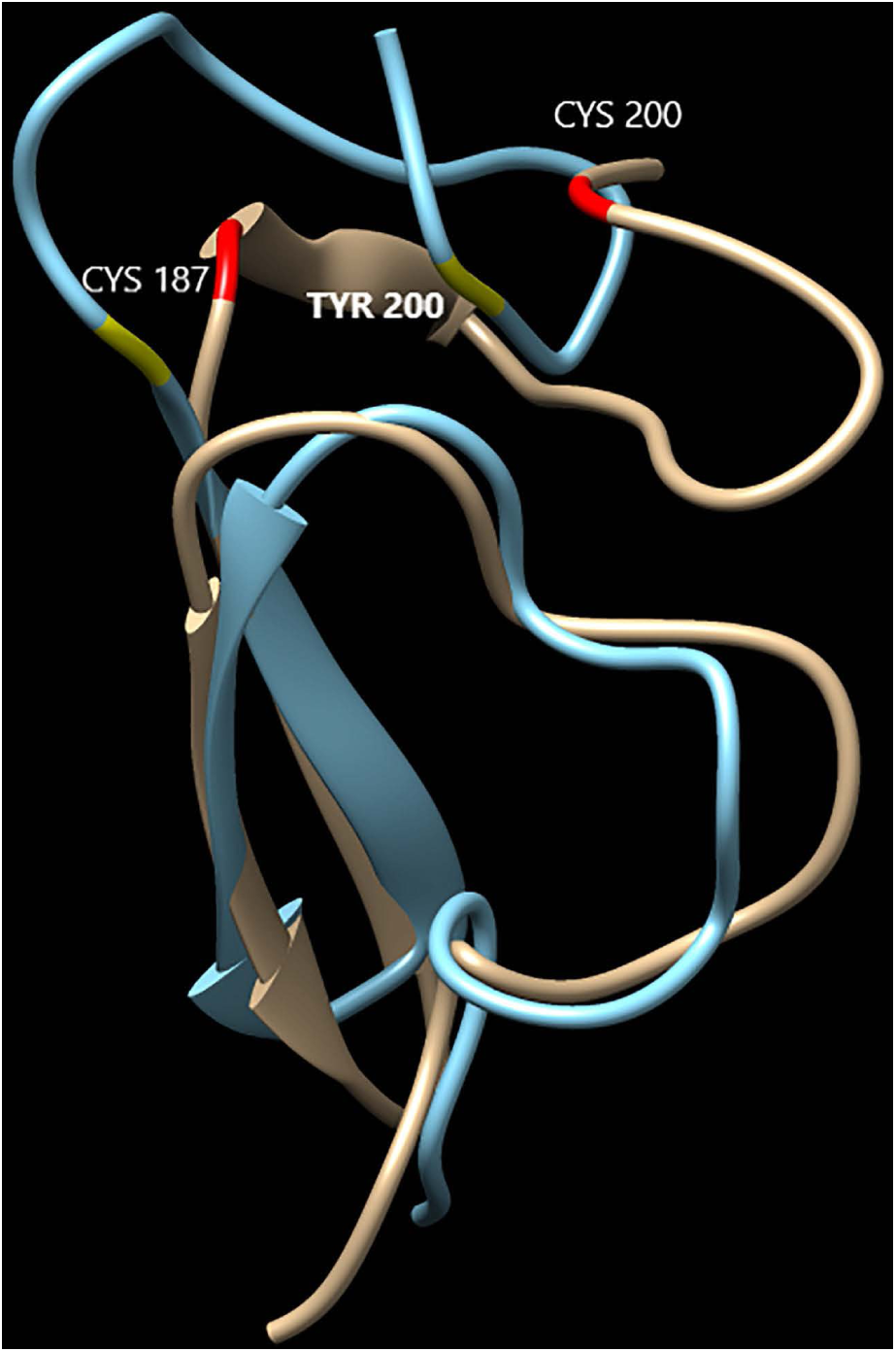
Protein interaction network from STRING database centered on *MEGF6* (A) and *ZBTB40* (B). Red nodes indicate the query protein and first shell of interactors, white nodes indicate second shell of interactors. Empty nodes indicate proteins of unknown 3-dimensional structure and filled nodes indicate 3-dimensional structure is known or predicted. Teal connectors indicate known interactions from curated databases, purple connectors indicate experimentally determined interactions, black connectors indicate co-expression, and yellow connectors indicate that interactions originate from text-mining.

### Analysis of zebrafish megf6a and megf6b gene expression and function

Our analyses indicated that a coding variant of *MEGF6* segregated with the occurrence of osteoporosis, that the variant was likely to be damaging to gene function, and that there was support for a functional link between the protein product of *MEGF6* and pathways known to contribute to osteoporosis. To uncover biological functions of *MEGF6*, we examined the consequences of complete loss of gene function in the zebrafish.

The zebrafish genome contains two *megf6* paralogues, *megf6a* and *megf6b*, which are orthologous to the single human *MEGF6* gene. Each orthologue has similar degree of identity (*megf6a* - 56% and *megf6b* – 54%) and similarity (*megf6a* - 68% and *megf6b* – 64%) to the human orthologue. The expression patterns of *megf6a* and *megf6b* were determined by whole-mount *in situ* hybridization (WISH) at 24, 36, and 48 hours post fertilization (hpf). At 24 hpf *megf6a* is broadly expressed in the head (Supplemental Figure 1A). At 36 hpf it is localized to the apical ectodermal ridge (AER) of the pectoral fin (Supplemental Figure 1B), the atrioventricular canal (Supplemental Figure 1C), and the notochord (Supplemental Figure 1D), and it is broadly expressed in the head. *megf6b* is expressed in the same domains as *megf6a* at 24 hpf, but it is also expressed in the developing somites (Supplemental Figure 1E). The expression pattern of *megf6b* is maintained in the fin fold and head at 36 hpf and is also expressed in the AER of the pectoral fin and in the notochord (Supplemental Figure 1F). Both *megf6a* and *megf6b* continue to be expressed in the notochord of 48 hpf larvae (data not shown). In the head of 48 hpf larvae, *megf6b* is expressed specifically in the mouth and pharyngeal region (Supplemental Figure 1G). These data indicate that *megf6a* and *megf6b* have mostly overlapping expression domains during the first 48 hpf of development and are expressed in regions of the embryo that give rise to cartilage and bone.

To generate zebrafish lacking both *megf6a* and *megf6b* gene function, we first generated a germline deletion of *megf6b*. Both *megf6a* and *megf6b* are large genes containing 34 and 38 exons, respectively. To make a large deletion in *megf6b*, we injected CRISPR/Cas9 ribonucleoprotein (RNP) complexes targeting exons 6 and 35 into one-cell stage embryos (26). We identified one allele, *z48* - a 51,860 bp deletion with a 53 bp insertion, which stably transmitted through the germline (Supplemental Figure 2). *megf6b* mutants are homozygous viable and the only overt phenotype is ectopic migration of a small number of melanocytes in the trunk and tail of 48 hpf larvae (Figure 4A-B’).

**Figure 4.**
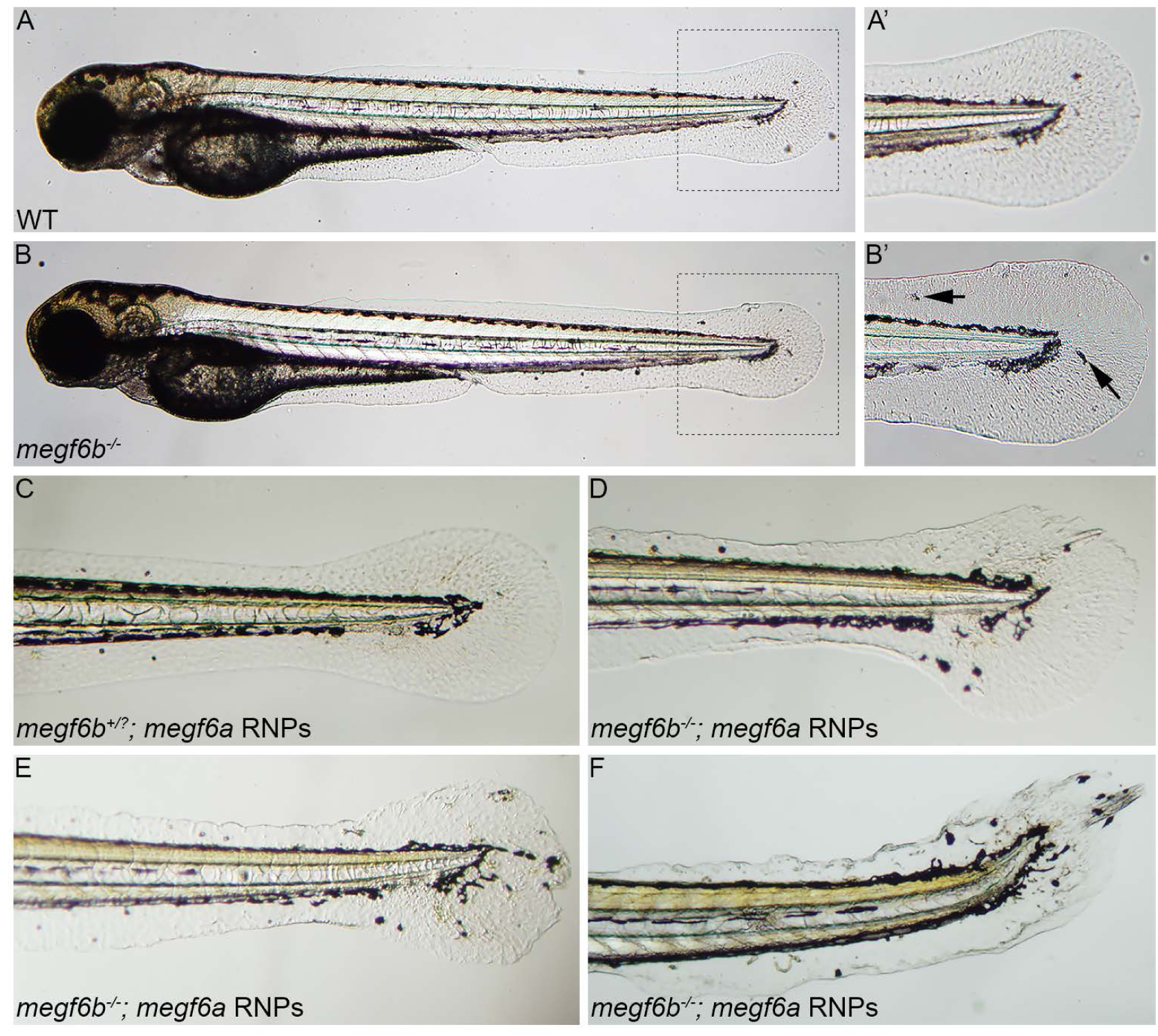
Larvae lacking *megf6a* and *megf6b* gene function have fin fold defects at 48 hpf. WT or *megf6b*^*-/+*^ larvae have no overt phenotypes (A and A’), while *megf6b*^-/-^ larvae have a few aberrant melanocytes (B’, arrow) present in the caudal fin area (B and B’). A’ and B’ are high magnification views of dashed boxes in A and B. (C-F) Injection of *megf6a* RNPs into larvae from an intercross between *megf6b*^*+/-*^ adults. Loss of *megf6a* gene function has no effect in a WT or *megf6b*^*+/-*^ background (C), while loss of *megf6a* gene function in *megf6b*^*-/-*^ larvae disrupted fin fold development and led to aberrant migration of melanocytes into the caudal fin area (D-F). All abnormal larvae were genotyped and determined to be *megf6b*^*-/-*^, indicating complete loss of *megf6b* function is required for the phenotype observed in the *megf6a* RNP-injected larvae (p < 0.0056, Fisher’s exact test with Bonferroni correction). Panels D-F depict the spectrum of aberrant phenotypes observed. Lateral views with anterior to the left.

Paralogous genes in zebrafish can have redundant biological functions (27). As *megf6a* and *megf6b* are both expressed in the fin fold, head, and notochord, we hypothesized they might have partially redundant functions. Therefore, we generated embryos lacking both genes by injection of *megf6b* mutant embryos with CRISPR-Cas9 RNPs that disrupt *megf6a* function. Our previous work demonstrated this CRISPR approach completely eliminates function of the targeted gene in the injected embryo and generates F0 embryos that faithfully recapitulate loss-of-function mutants (26). We injected CRISPR-Cas9 RNPs targeting exons 6 and 28 of *megf6a* into one-cell stage embryos generated from an intercross between *megf6b*^+/-^ adults, providing both control wildtype and *megf6b* mutant embryos that lacked *megf6a* function.

Whereas three-quarters of the intercross embryos appeared phenotypically wildtype following CRIPSR-Cas9 mutagenesis of *megf6a* (Figure 4C), roughly one-quarter of the larvae (21.7%, (n =115)) exhibited a severe disruption of the fin fold with aberrant migration of melanocytes into the fin fold area (Figure 4 D-F). The aberrant embryos were genotyped, and all were found to be homozygous for the *megf6b* mutation, indicating that mutagenesis of *megf6a* only has measurable effect on fin fold development in the absence of *megf6b* function (p = 0.0056). These results indicate *megf6a* and *megf6b* have redundant functions during embryogenesis: loss of either gene had minimal effect on development whereas larvae lacking both *megf6a* and *megf6b* gene functions had severe defects on fin development.

Cartilage and bone form in an anterioposterior wave of development in the zebrafish larva, so that at any single time point posterior structures are developmentally younger than anterior ones. Therefore, to examine cartilage and bone formation in the head and in the vertebral column, we analyzed mutant larvae at two developmentally distinct time points, 10 and 14 days post fertilization (dpf). To determine if *megf6a* and *megf6b* have a function in cartilage and bone development in the head, we analyzed 10 dpf larvae that developed from *megf6b*^+/-^ intercross eggs injected with *megf6a* RNPs. Cartilage and bone formation were visualized after staining larvae with Alcian Blue (cartilage) and Alizarin Red (bone) (28). Injection of *megf6a* RNPs into WT or *megf6b*^+/-^ larvae had no measurable effect on bone and cartilage formation (Figure 5A and 5A’). However, twenty percent of the injected intercross larvae (n = 110) exhibited consistent delays in jaw development, including formation of the pharyngeal arches and development of both bones (5th ceratobranchyal (5CB) or cleithrum (CL)) and cartilaginous structures of the jaw (Figure 5B and 5B’). Genotyping revealed that all the injected embryos with jaw defects were *megf6b*^*-/-*^, indicating that loss of both *megf6* genes was required (p = 0.0062) to produce significant delays in jaw development.

**Figure 5.**
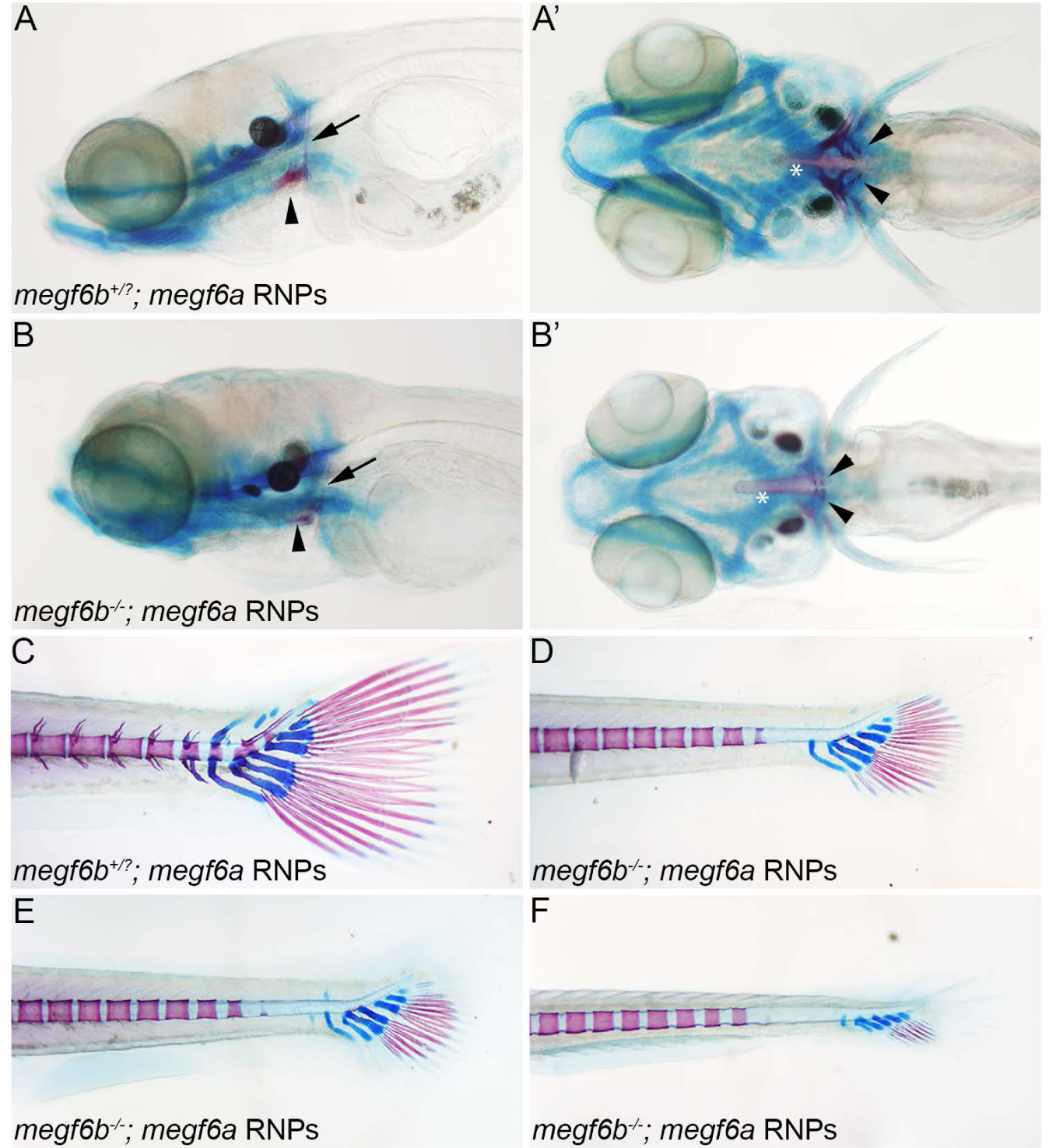
Cartilage and bone formation is delayed in larvae lacking both *megf6a* and *megf6b* gene function. Progeny of an *megf6b*^*-/+*^ intercross were injected with *megf6a* RNPs. (A-B’) Cartilage and bone formation in the heads of RNP-injected WT, *megf6b*^*+/?*^ (A and A’), or *megf6b*^*-/-*^ (B and B’) larvae were analyzed at 10 dpf. Arrowhead indicates the 5th ceratobranchyal (5CB) and arrow indicates the cleithrum (CL). Twenty-two of 110 larvae exhibited delayed formation of jaw bones and pharyngeal arches, and all of these were *megf6b*^*-/-*^, indicating loss of both genes is required for the developmental defect (p < 0.0062, Fisher’s exact test with Bonferroni correction). A and B, lateral view and A’ and B’, ventral view. Asterisk indicates the notochord. (C-F) Cartilage and bone formation in in the tails of RNP-injected WT, *megf6b*^*+/?*^, or *megf6*^*-/-*^ larvae were analyzed at14 dpf. Mineralization in the vertebra and formation of cartilaginous structures of the caudal fin are significantly delayed in larvae lacking *megf6a* and *megf6b* function (17.5%, n=80) (D-F) compared to WT or *megf6b*^*+/?*^ (C) larvae injected with *megf6a* RNPs. All abnormal larvae were genotyped and determined to be *megf6b*^*-/-*^ indicating loss of both genes is required for the delayed phenotype (p < 0.007, Fisher’s exact test with Bonferroni correction). Panels D-F depict the spectrum of phenotypes observed when *megf6a* RNPs are injected into *megf6b*^*-/-*^ embryos. C-F, lateral views with anterior to the left.

Given that *megf6a* and *megf6b* have overlapping expression domains in the notochord (the embryonic version of the vertebral column) and that the two genes have overlapping functions in fin fold development (Figure 4D-F), we analyzed the roles of the genes in bone and cartilage formation in the vertebral column and caudal fin of 14 dpf larvae. Progeny of a *megf6b*^+/-^ intercross mating were injected with *megf6a* RNPs. Most larvae exhibited normal bone and cartilage formation in the vertebral column and caudal fin (Figure 5C). However, 17.5% of the larvae (n=80) exhibited a severe delay in the ossification of the posterior vertebrae and a delay in the formation of the cartilaginous structures and bony rays of the caudal fin (Figure 5D-F). All the abnormal larvae proved to be *megf6b*^-/-^ upon genotyping, indicating that only embryos lacking both *megf6b* and *megf6a* gene functions had an altered phenotype (p < 0.007). The functions of both genes are required for proper formation of both cartilaginous and bony structures in the zebrafish larvae.

To determine if zebrafish lacking *megf6a* and *megf6b* function have an adult phenotype, we separated 48 hpf larvae that developed from *megf6b*^+/-^ intercross eggs injected with *megf6a* RNPs (Figure 4) into two groups of 25 animals, those with normal fin folds (*megf6b*^*-/?*^) and those with fin fold defects (*megf6b*^*-/-*^). Despite the developmental delay observed in doubly mutant larvae, 100% of the individuals of each group developed to adulthood. At 4 months of age, we measured the weight (g) and length (cm) of individual male and female adult zebrafish. The males in the abnormal fin fold group weighed less (mean ± SD - 0.4098g ± 0.014g) than the males with normal fin folds (0.4556g ± 0.014g), while there was no significant difference in length between the two groups (abnormal fin folds – 3.671cm ± 0.032cm vs normal fin folds – 3.721cm ± 0.035cm). In contrast to the males, females in the abnormal fin fold group weighed more (abnormal fin folds - 0.6616g ± 0.021g vs normal fin folds 0.5332g and ± 0.020g) and were longer (abnormal fin folds – 3.971cm ± 0.035cm vs normal fin folds – 3.809cm ± 0.034cm) than females with normal fin folds (Supplemental Figure 3).

## DISCUSSION

Whole exome sequence data from pairs of related osteoporosis-affected individuals from 10 extended high-risk pedigrees were analyzed to identify a small set of rare, IBD-shared variants representing excellent candidates responsible for osteoporosis predisposition in the pedigrees studied. A rare *MEGF6* coding variant segregated with osteoporosis in the pedigree in which it was identified. *In silico* analyses of the predicted consequences of the amino acid substitution indicated the variant is likely to be damaging to normal protein function. STRING analysis indicated that MEGF6 may interact with proteins linked to osteoporosis. Loss-of-function analyses in the zebrafish revealed *megf6* gene function contributes to cartilage and bone formation. Together these support the interpretation that the very rare variant in *MEGF6* predisposes to osteoporosis.

The design of the current study focused on exome sequencing of pairs of related affected individuals selected from extended high-risk pedigrees. Selection of subjects focused on affected cousins, which has been shown to be ideal when selecting subjects to represent dense pedigrees (29). The main benefit of the affected-relative-pairs strategy is efficiency with respect to data capture, but at the potential cost of missing evidence of disease variants that may have been detectable in a design that sequences all pedigree members. Here the strategy was sufficiently robust to lead to identification of at least one variant in ten pedigrees. The reliance on SGS analysis to delineate regions of long-range sharing between target pair members provided a productive filter of potential candidate variants. The use of well-sampled high-risk pedigrees has the added advantage of being able to invoke a test of segregation among additional affected family members to gain evidence of linkage to disease, which was shown here to be a particularly useful strategy for confirmation of the very rare variant rs755467862.

Consistent with our current finding of an association between *MEGF6* and osteoporosis, previous studies had observed an association between low BMD or osteoporosis and the chromosomal region containing the *MEGF6* locus (30, 31). A LOD score of +2.74 to 1p36.32 was reported in a resource of 1,270 subjects in 324 osteoporosis pedigrees (30). Independently, a LOD score of +3.07 to 1p36.3 was reported in a single Belgian pedigree with 34 subjects (31). In the latter study, it was reported that exon sequencing of *MEGF6* did not identify any coding variants among pedigree members.

*In silico* modeling of the WT and variant MEGF6 indicate that the Cys200Tyr may result in collapse of a loop that likely eliminates a calcium binding site in MEGF6, which may decrease, or even eliminate, the functionality of the protein. Similar arguments have been made by observation of mutations of EGF-like domains in other proteins. For instance, more than 60% of the mutations causing Marfan syndrome occur within fibrillin-1 cbEGF-like domains, emphasizing that correct folding of these domains is critical for molecular function (32). Mutations in cbEGF-like domains that affect cysteine residues are likely to alter disulfide bond formation, which may disrupt correct folding (32). Furthermore, mutations affecting residues in the calcium binding consensus sequence reduce calcium binding affinity, which may lead to structural destabilization (32).

The biological roles of MEGF6 are not well understood. MEGF6 appears to be a secreted calcium binding protein that regulates cell migration through TGF-β/SMAD mediated epithelial-mesenchymal transition (33). Depletion of *MEGF6* caused reduction of TGF-β1 expression and phosphorylation of SMAD2 and SMAD3, whereas *MEGF6* expression positively correlated with that of TGF-β1 expression, suggesting a role for *MEGF6* in TGF-β/SMAD signaling. In osteoporosis, TGF-β is proposed to regulate osteoclastogenesis through activation of osteoprotegerin (OPG) (34). OPG has been shown to inhibit the activity and survival of osteoclasts in vitro and bone resorption *in vivo* (23). It is possible that reduced *MEGF6* activity may alter TGF-β signaling and thus OPG expression, thereby contributing to osteoporosis. Furthermore, we identified another potential link between *MEGF6* and osteoporosis through a ZBTB40 -WNT16 interaction using the STRING database. Variants in *ZBTB40* and *WNT16* have been previously associated with osteoporosis (25, 35) and osteoblast-derived *WNT16* inhibits osteoclastogenesis to prevent bone fragility (36). Further studies are needed to determine if *MEGF6* regulates TGF-β signaling *in vivo* and if it interacts with *ZBTB40* or *WNT16*.

Our functional analyses demonstrate that the zebrafish orthologues of human *MEGF6, megf6a* and *megf6b*, are together required for proper development and mineralization of both cartilage and bone in the head, vertebrae, and caudal fin. In the head of 10 dpf larvae, we observed a delay in the formation of the cartilaginous pharyngeal arches and a delay in mineralization of the 5th ceratobranchyal (5CB) and the cleithrum (CL) bones, while there was no defect in the initial mineralization of the notochord (Figure 5A-B’). At 14 dpf there was a delay in the mineralization of the vertebral column and bones of the caudal fin, and a delay in the formation of cartilaginous structures of the caudal fin (Figure 5C-F). The mineralization of the 5CB, CL, and notochord (vertebral column) represent distinct modes of bone ossification: endochondral, membranous, and perichordal bone ossification, respectively. Given there is a delay in mineralization of all of these structures, *megf6a* and *megf6b* have important functions in the distinct modes of bone ossification.

The bone phenotype observed in larvae lacking *megf6a* and *megf6b* function is strikingly similar to several previously reported zebrafish mutant phenotypes. A mutation in the *macrophage-stimulating protein* (*msp*) gene, or morpholino knockdown of its receptor, *ron-2*, display similar defects in the 5CB bone (37). The *msp* mutant phenotype can be rescued by exogenous calcium supplementation to the media, indicating that Ron-2 and Msp are necessary for bone mineralization and calcium homeostasis (37). The zebrafish *Chihuahua* (*Chi*) mutant disrupts *col1a1a* and is a model for classical osteogenesis imperfecta. Heterozygous *Chi* mutants have the same defects in the 5CB and CL bones and a delay in vertebral mineralization as are conspicuous in larvae with disrupted *mefg6a* and *megf6b* function (38). The zebrafish *frilly fins* (*frf*) mutant disrupts the *bmp1a* gene and is phenotypically similar to the *microwaved* mutant, which carries another mutant allele of *col1a1a* (39). BMP1 has a known function in the proteolytic processing of procollagen I C-propeptide to generate mature collagen type I (40), which may account for the phenotype similarities between the mutants. The *frf* mutant has a delay in the mineralization of the vertebrae reminiscent of larvae without *megf6a* and *megf6b* gene function (39). Given the similarity of these mutant phenotypes and with the perspective that *MEGF6* is a calcium binding protein, *MEGF6* may have a role in regulating calcium homeostasis and collagen processing during embryonic development.

A *Megf6*-deficient mouse was recently reported with no discernable phenotype (41). The apparent discrepancy between the mouse and zebrafish mutants may lay in the phenotypic analyses performed. There was no detailed histological analysis of embryonic skeletal development in the mouse, which may have overlooked subtle defects, and BMD was not analyzed in the adult. Skeletal phenotypes may be uncovered upon a more detailed analysis, since *Megf6* is expressed in the mouse musculoskeletal system during development (42), during osteoblast differentiation, and in mature bone (43). Furthermore, as the mouse mutation involved a coding alteration that resulted in no detectable *Megf6* transcript (41), the mutant transcript likely underwent non-sense mediated decay and may have triggered genetic compensation mechanisms that masked the null phenotype (44, 45).

Despite the dramatic developmental fin fold phenotype of *megf6b*^*-/-*^ larvae injected with *megf6a* CRISPR/Cas9 reagents, adult zebrafish do not exhibit severe phenotypes (e.g., kyphosis), although there is a significant difference in the weight and length of these zebrafish. The adult males with abnormal fin folds weigh less than controls with normal fin folds without a reduction in overall length, which may reflect a loss in bone density. In contrast to the adult males, adult females with abnormal fin folds weigh more and are longer than controls with normal fin folds (Supplemental Figure 3). This phenotype may be due to factors unique to the teleost lineage, including sex differences and compensatory growth mechanisms. Zebrafish grow in size throughout their lifetime and have prodigious regenerative capacities (46, 47), which may contribute to the mild adult phenotype and the observed phenotypic differences between males and females. Further characterization of the zebrafish *megf6a;6b* mutant phenotype in aged adults and generation of zebrafish with the human disease allele (48) will provide insight into the genetic mechanisms of osteoporosis and may yield new targets for therapeutic intervention.

In sum, evidence of segregation, protein folding prediction, and *in vivo* functional evaluation support a role for *MEGF6* in predisposition to development of osteoporosis.

## MATERIALS AND METHODS

### Utah osteoporosis pedigrees

Osteoporosis cases with a self-reported family history were ascertained and recruited from clinics associated with Intermountain Healthcare, the largest Utah health care provider, and local advertising in health care clinics; relatives of these cases were also recruited. All recruited individuals were offered a BMD scan for determination of osteoporosis phenotype. BMD was measured on a Hologic QDR 4500A fan-beam dual X-ray absorptiometry (DXA) bone densitometer by a single technician (49, 50). Quality control for the DXA was performed daily on a spine phantom and all scans were reviewed and evaluated by a single experienced physician. BMD was measured at the hip and spine (or at the wrist if the hip or spine could not be imaged). In addition to the anteroposterior (AP) view of the spine, lateral measurements were taken. Scans were excluded if their interpretation was confounded by poor positioning or abnormal anatomy.

A total of 1,871 individuals were consented, phenotyped, and sampled in 276 multi-generational pedigrees with 4–46 sampled individuals per pedigree. In addition to BMD measurements, a single physician reviewed a subject medical history questionnaire, and available medical records to assign a phenotype of osteoporosis, osteopenia, unaffected, or unknown. Ten pedigrees were selected for sequencing based on a physician diagnosis of osteoporosis in a pair of (approximately) cousins (29), and having additional sampled affected relatives for tests of segregation. This study was approved by the Institutional Review Boards of Intermountain Healthcare and the University of Utah.

### Utah genomic data

Whole exome sequencing was performed on the 10 pairs of affected cousins at the Huntsman Cancer Institute’s Genomics Core facility. DNA libraries were prepared from 3 micrograms of DNA using the Agilent SureSelect XT Human All Exon + UTR (v5) capture kit. Samples were run on the Illumina HiSeq 2000 instrument. Reads were mapped to the human genome GRCh37 reference using BWA-mem (51) for alignment and variants were called using Genome Analysis Toolkit (GATK) (52) software following Broad Institute Best Practices Guidelines. Exome capture resulted in an average of 85% of target bases being covered by greater than 10x coverage across the genome. Variants occurring outside the exon capture kit intended area of coverage were removed. Variants were annotated with Annovar, which contains predicted pathogenicity scores from 10 in-silico functional prediction algorithms (11). Samples were also genotyped on the Illumina OmniExpress high density SNP array (720,000 SNPs).

### Candidate variant prioritization

Variants were filtered in four phases to achieve a small set of highly probable candidates. In the first phase, variants with population minor allele frequency (MAF) <0.005 that were observed in both index cases from a pedigree (affected cousins) were retained. In the second phase of variant filtration, the genomes of index pairs were restricted to regions exhibiting evidence of IBD transmission, or shared common ancestry. Focus on these IBD regions between sequenced cousin-pairs provides a useful filter that can rule out variants that are shared but were not inherited from a common ancestor and are inconsistent with a dominant mode of inheritance.

SNP genotypes were used to estimate regions of long shared haplotypes between related sequenced cases using SGS (8). SGS analysis identifies the set of contiguous markers at which genotyped individuals could share alleles; long runs of such markers indicate likely regions of IBD sharing from a common ancestor (53). Genomic regions were considered as IBD for a sequenced relative pair if the normalized shared-segment length was ≥20 standard deviations from the mean shared-segment length (54). This selection criteria was based on the expected amount of IBD sharing that should occur for a pair of cousins (∼12.5%), and provided an equivalent reduction to the genomic search space. The third phase of variant filtration required that the gene had at least one published study connecting the gene with osteoporosis through some genetic study design, identified via PubMed search (55) with search terms ‘osteoporosis’ and gene name or alias provided by OMIM (Online Mendelian Inheritance in Man, OMIM®. McKusick-Nathans Institute of Genetic Medicine, Johns Hopkins University (Baltimore, MD), January 17, 2018: https://omim.org/). In the final filtering step, variants that were scored as ‘damaging’ by at least 8 of 10 in-silico pathogenicity prediction algorithms included in ANNOVAR variant annotations were retained.

### Confirmation of candidate variants via segregation to affected relatives

The 10 pedigrees of the sequenced relative pairs had a total of 68 additional sampled relatives, ranging from 4 to 15 per family. To confirm segregation of the candidate predisposition variants in the pedigrees, each variant was assayed in the sampled relatives of sequenced cases using custom Taqman assays (Applied Biosystems, Carlsbad, CA, USA) according to the manufacturer’s specifications on a Bio-Rad CFX96 Real-Time PCR instrument. Evidence of segregation to additional affected relatives was evaluated with RVsharing software (9) that provides the probability of the observed configuration of sharing in a pedigree, expressed as an exact probability, assuming the shared variant is rare (<1%) and entered the pedigree only once.

### Pathway analysis and protein structure prediction

The STRING database tool (21) with default settings was used to query a large number of known protein interactions concerning the candidate gene. Protein structure prediction software I-TASSER using full homology modeling was used to model the structure of critical domains assuming either wild-type or variant amino acid sequences (19, 56-58). Predicted structures were visualized and analyzed using Chimera software (59).

### Zebrafish husbandry

Zebrafish (Danio rerio) were maintained in accordance with approved institutional protocols at the University of Utah. Adult zebrafish were maintained under standard conditions (60) and kept on a light-dark cycle of 14 hours in light and 10 hours in dark at 27°C. The Tu strain was used in all experiments.

### megf6a and megf6b Mutant Generation

Mutations were generated with CRISPR-Cas9 reagents as described in (26). gRNA target sequences were as follows: *megf6a* exon 6 - CGCTGTCAGCATGGTGTTCT(TGG) and exon 28 - GCTCGCAGTGACCCTCTCAC(AGG); and *megf6b* exon 6 - AGCACACATGTGTGAACACT(AGG) and exon 35 -TCCTGTATCCGGATGACAGG(AGG). The PAM sequence is indicated in parentheses. Target-specific Alt-R® crRNA and common Alt-R® tracrRNA were synthesized by IDT and dissolved in duplex buffer (IDT) as a 100μM stock solution. Equal volumes of the Alt-R® crRNA and Alt-R® tracrRNA stock solutions were mixed together and annealed in PCR machine using the following settings: 95°C, 5 min; cool at 0.1°C/sec to 25°C; 25°C, 5 min; 4°C. Cas9 protein (Alt-R® S.p. Cas9 nuclease, v.3, IDT, dissolved in 20mM HEPES-NaOH (pH 7.5), 350mM KCl, 20% glycerol) and crRNA:tracrRNA duplex mixed to generate a 5μM gRNA:Cas9 RNP complex (referred to as RNPs). Prior to microinjection, the RNP complex solution was incubated at 37°C, 5 min and then placed at room temperature. Approximately one nano-liter of 5μM RNP complex was injected into the cytoplasm of one-cell stage embryos.

### Genomic DNA extraction and genotyping

Genomic DNA was extracted from individual embryos at 24 hours post fertilization (hpf). Dechorionated embryos were incubated in 50 ul 50 mM NaOH at 95°C, 20 min. 1/10 volume of 1 M Tris-HCl (pH 8.0) was added to neutralize. Genome sequences containing CRISPR/Cas9 target sites were amplified with pairs of primers: *megf6a* ex6 HRMA F1 – GGAGATTGTCAATACCTGTGACT and *megf6a* ex6 HRMA R1 – AGGTCTTCGGCGAGCTGATA; *megf6a* ex28 HRMA F1 – GGTCAGGACTGTGCTGGAGTG and *megf6a* ex28 HRMA R1 – CCCACAGTCCCGTCCACGC; *megf6b* ex6 HRMA F1 – ATATTGACGAGTGCCAGGTTCAT and *megf6b* ex6 HRMA R1 – ATGCAGTCGAGAGCCGGCGTTA; *megf6b* ex35 HRMA F1 – GCTCTGGGTGTCAGCAGCAGT and *megf6b* ex35 HRMA R1 – GCTGACAGCGCGGGCCGT. To determine if individual gRNA:Cas9 RNPs produced mutations at the desired target sites, high resolution melt analysis (HRMA) was performed on DNA isolated from 8 individual 24 hpf gRNA:Cas9 RNP-injected embryos using KAPA HRM FAST PCR Master Mix (61). To generate the *megf6b* deletion allele, a mixture of gRNA:Cas9 RNPs targeting exons 6 and 35 was injected into the cytoplasm of one-cell stage embryos. To detect deletion events, PCR was performed with *megf6b* ex6 HRMA F1 and *megf6b* ex35 HRMA R1 primers on DNA isolated from 8 individual 24 hpf G0 gRNA:Cas9 RNP injected embryos using KAPA 2G FAST PCR Master Mix. To remove *megf6a* gene function in G0 embryos, a mixture of gRNA:Cas9 RNPs targeting exons 6 and 28 was injected into the cytoplasm of one-cell stage embryos.

### In Situ Hybridization

Whole mount in situ hybridization (WISH) was performed on embryos as described in (62). cDNA fragments used to generate riboprobes probes were amplified by RT-PCR using the following primers: *megf6a* ISH F – TGGCACCTGCAGCTGCCC, and *megf6a* ISH R – TCCAGCCGTTCAGACACGTGCA, or *megf6b* ISH F – TGAACAGACGTGTCCGCAGGG, and *megf6b* ISH R – TCACACTCGCACAGCAGAGAGC using KAPA 2G FAST Master Mix, sub-cloned into pGEM-T Easy, and subjected to Sanger sequencing for verification.

### Cartilage and Bone Staining

Ten and 14 dpf zebrafish larvae were anesthetized with Tricaine methanesulfonate (ethyl 3-aminobenzoate methanesulfonate) and processed as described in (28) and https://wiki.zfin.org/pages/viewpage.action?pageId=13107375 with the following modifications. Larvae were fixed in 2% paraformaldehyde for 1 hour, washed for 10 minutes in 50% EtOH, and then transferred to a solution containing 0.01% Alizarin Red and 0.04% Alcian Blue for 24 hours. Larvae were washed in 80% EtOH/10mM MgCl2 for 60 minutes, 50% EtOH for 30 minutes, 25% EtOH for 30 minutes, bleached in 3% H2O2/0.5% KOH for 15 minutes, washed in 2X 25% glycerol/0.1% KOH and then transferred to 50% glycerol/0.1% KOH for imaging.

### Length and Weight Measurements

One-cell stage embryos from an intercross between *megf6b*^*+/-*^ adults were injected with *megf6a* RNPs and larvae were sorted at 48 hpf into two groups of 25 animals, those with normal fin folds (*megf6b*^*-/?*^) and those with fin fold defects (*megf6b*^*-*/*-*^). Animals were raised to 4 months of age. At 4 months of age adults were anesthetized using Tricaine methanesulfonate and the length (cm) of each zebrafish was determined by measuring from the anterior most portion of the head to the tip of the tail. Zebrafish were then weighed and allowed to recover from anesthesia.

### Statistical analysis

One-cell stage embryos from an intercross between *megf6b*^*+/-*^ adults were injected with *megf6a* RNPs and larvae were sorted at 48 hpf based on whether or not they displayed the mutant fin fold phenotype. Embryos were then genotyped as describe above. For the cartilage and bone analysis, embryos were stained, sorted based on phenotype, and then genotyped. To determine whether phenotypically defective larvae were significantly enriched for the *megf6b*^*-/-*^ genotype, Fisher’s exact test with Bonferroni correction was used to determine statistical significance. A Student’s t-test was used to determine statistical significance for the weight and length comparisons.

## ACKNOWLEDGEMENTS

The samples used for this project originated from a collaboration between Myriad Genetics, Intermountain Healthcare, and the University of Utah. We are especially grateful to the families who participated in this study. University of Utah Core Facilities provided support for oligonucleotide synthesis, DNA sequencing, and maintenance of zebrafish.

## FIGURE LEGENDS

**Supplemental Figure 1**. *megf6a* and *megf6b* have both overlapping and unique expression domains. Whole mount *in situ* hybridization of *megf6a* (A-D) and *megf6b* (E-G) at 24 hpf (A and E), 36 hpf (B, C, D, and F), and 48 hpf (G). *megf6a* and *megf6b* are expressed in the fin fold (arrow in A) and in the head of 24 hpf embryos (A and E), while only *megf6b* is expressed in somitic tissue (arrow in E). At 36 hpf *megf6a* and *megf6b* are expressed in the fin fold (D and F), AER (arrowhead in B and F), atrioventricular canal (arrowhead in C), and notochord (arrow in D and F), and head. *megf6b* is expressed in the mouth (arrow in G) and pharyngeal region (arrowheads in G) in 48 hpf larvae. Lateral views with anterior to the left.

**Supplemental Figure 2**. Generation of a heritable *megf6b* deletion mutant. (A) RNPs were designed to target sites (lightning bolts) within exon 6 (ex 6 gRNA) and exon 35 (ex 35 gRNA) of *megf6b.* Primer pairs used to amplify junction fragments joining target sites are indicated (red arrows). (B) Detailed view of the ex 6 gRNA and ex 35 gRNA sites with gRNA target sequence in red, PAM site in blue, and Cas9 cleavage site indicated by lightning bolts. *megf6bΔ ex 6/35* represents a conceptual precise joining of sequences flanking the presumed cleavage sites in target sites ex 6 and ex 35. *megf6b*^*z48*^ represents the deletion allele recovered in the germline. This allele contains a 51,860 bp deletion with a 53 bp insertion.

**Supplemental Figure 3**. Weight (g) and length (cm) of adult zebrafish lacking *megf6a* and *megf6b* function. The weight (A) and length (B) of individual male and female 4 month old zebrafish generated from *megf6b*^*+/-*^ intercross eggs injected with *megf6a* RNPs. Circles indicate embryos identified at 48 hpf with normal fin folds (*megf6b*^*+/?*^) and squares indicate those with fin fold defects (*megf6b*^*-/-*^). Males with abnormal fin folds weighed less than controls with normal fin folds (A), while there was no difference in length between the two groups (B). Females with abnormal fin folds weighed more and were longer when compared to controls with normal fin folds (A and B). n = 14 males and n = 11 females with normal fin folds and n= 14 males and n = 7 females with abnormal fin folds. A Student’s t-test was used to determine statistical significance. Data are represented as means ± SD. *p ≤ 0.05; **p ≤ 0.01; ***p ≤ 0.001.

