## Supplementary figures and images for "A role for the *MEGF6* gene in predisposition to osteoporosis"

### Supplemental Figure

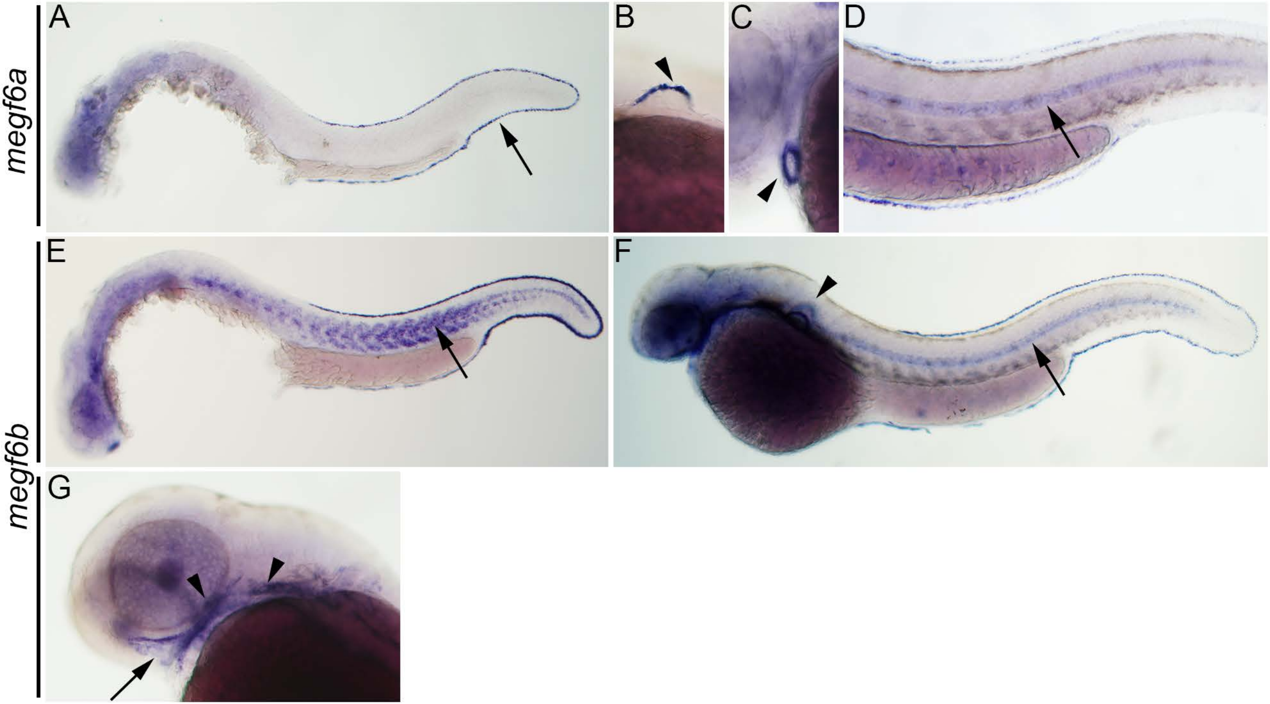

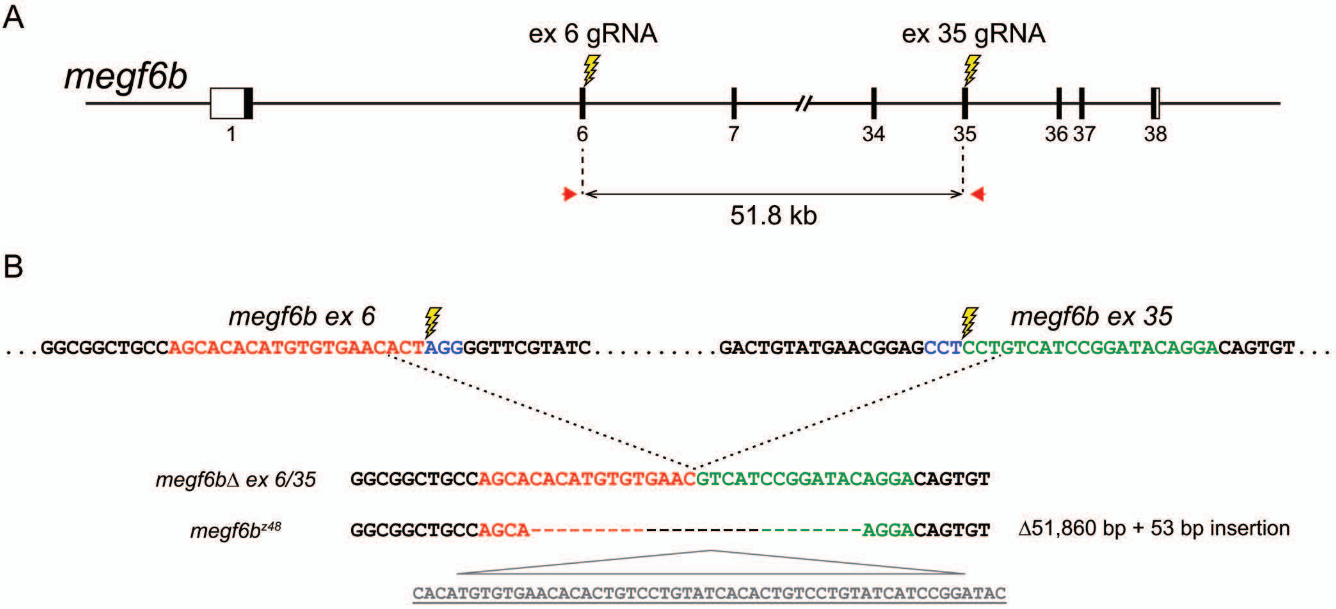

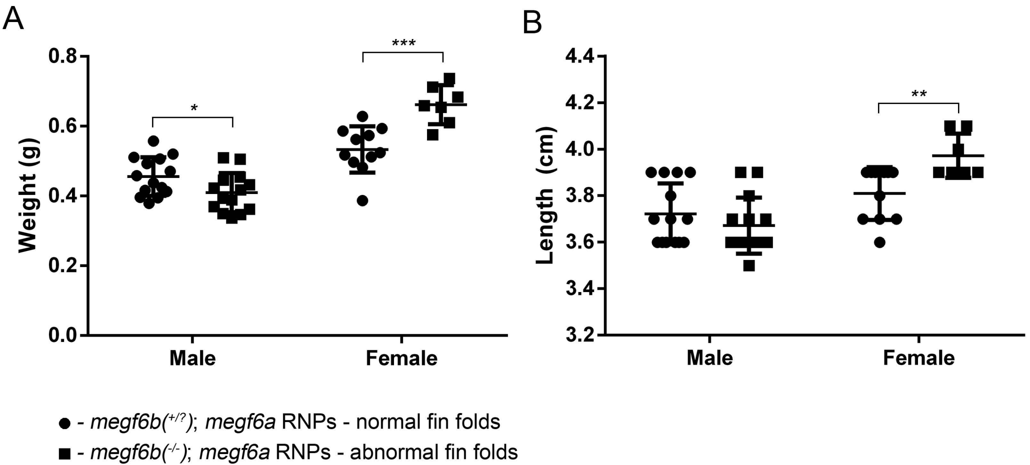
